# AgRP Neuron Activity Predicts and Tracks the Glycemic Response to Oral Glucose

**DOI:** 10.64898/2026.04.01.715678

**Authors:** Micaela Glat, Anna J. Bowen, Yang Gou, Elizabeth Giering, Jarrad M. Scarlett, Gregory J. Morton, Michael W. Schwartz

## Abstract

Hypothalamic AgRP neurons are uniquely responsive to nutritional cues and play an important role in fuel homeostasis. To investigate the temporal relationship between the activity of these neurons and the glycemic response to an oral glucose load, we simultaneously monitored AgRP neuron activity (by fiber photometry in AgRP-IRES-cre mice) and the arterial glucose level, both before and after oral gavage (OG) of either water or glucose (0.5-2.5 g/kg). We report that the AgRP neuron response to an OG glucose load can be subdivided into two functionally distinct phases – one that begins prior to glucose delivery and a second that extends from peak inhibition through the return towards baseline. The ‘first phase’ appears to be anticipatory in nature and is also predictive of subsequent changes in glycemia, suggesting a role in the handling of an oral glucose load. To analyze the relationship between the second phase response and changes of glycemia, we employed a model that allows residual activity to be removed subsequent to the ‘first phase’ component. This analysis reveals that unlike the first phase, the degree of residual inhibition – the second phase – tracks the glycemic response. Moreover, this response is temporally aligned with the blood glucose (BG) rate of change (which is predictive of future BG levels), with AgRP neurons lagging BG rate of change by ~5 minutes. We conclude that the AgRP neuron response to an oral glucose challenge consists of two distinct phases, each with its own determinants and metabolic implications: an initial anticipatory component that is predictive of the subsequent glycemic response, and a second phase that tracks the rate of BG change.

## 1. Introduction

The glycemic excursion following an oral glucose load is maintained within narrow physiological limits by dynamic and highly integrated interactions across central and peripheral tissues throughout the body [1]. Contributing factors include the rate of gastric emptying, the secretion of incretin peptides (such as glucagon-like peptide-1 (GLP-1) and gastric inhibitory polypeptide (GIP)) from intestinal enteroendocrine cells, and the interaction between insulin secretion and tissue responsiveness to insulin [2–4]. In addition, ‘gut-brain’ signals transmitted by vagal and somatosensory afferent neurons as well as incretin peptides convey changes of nutritional state from the gastrointestinal (GI) tract to the brain, which in turn mounts autonomic and neuroendocrine responses that shape glucose handling [5,6]. Importantly, a subset of these responses is initiated before nutrient ingestion, in anticipation of the coming nutrient load [4].

Neurons located in the hypothalamus play key roles in this control system. Among the best-studied of these are AgRP neurons, situated in the hypothalamic arcuate nucleus (ARC) [7]. These neurons are activated by fasting [8], uncontrolled diabetes [9,10], and other fuel-depleted states. This activation response is rapidly reversed by nutrient presentation [11–13] – not only by the consumption of food, but by infusion of macronutrients directly into the GI tract [7] or even just the sight or smell of food [7,13,14].

These observations highlight how AgRP neuron activity is regulated not only by changing nutritional status, but also by the anticipation that nutritional status will soon change. Relevant nutrient-related cues implicated in this regulation include gut-brain signals, hormones such as insulin, leptin and ghrelin, and circulating nutrients such as glucose [12,15,16]. Given that AgRP neurons are activated in most forms of diabetes [7,17–19], the observation that hyperglycemia can be normalized by silencing them in a mouse model of type 2 diabetes [17] highlights the important role these neurons can play in obesity-associated glucose metabolic impairment.

Such observations add salience to the question of whether and how AgRP neuron activity and changing BG levels are linked to one another. To address this question, we simultaneously monitored AgRP neuron activity (by fiber photometry) and the arterial blood glucose (BG) level (by continuous glucose monitoring (CGM)) in 5-hour fasted mice. Data were collected both prior to and after oral gavage (OG) of either water or one of four doses of glucose (0.5-2.5 g/kg), and computational modeling was employed to quantify these relationships.

We report that the AgRP neuron response to an oral glucose load can be subdivided into two distinct phases. The first of these is a rapid inhibitory phase that begins prior to glucose delivery, suggestive of an anticipatory response, and is predictive of the subsequent dose-independent glycemic excursion. Following a brief transitional period during which peak neuronal inhibition occurs, the ‘second phase’ of the response (during which AgRP neuron activity returns towards its baseline value) occurs and is temporally linked to the glycemic excursion – specifically, to the BG *rate of change*. Collectively, these observations highlight a multiphasic and temporally structured relationship between AgRP neuron activity and the glycemic response to an oral glucose load, consistent with the known multimodal responsivity of these neurons [13,14].

## 2. Results

### 2.1

To identify relationships between AgRP neuron activity and the blood glucose level following an oral glucose load, we combined fiber photometry (FP) with continuous arterial glucose monitoring (CGM). AgRP neuron-specific expression of a Cre-dependent AAV encoding the calcium reporter GCaMP6s (**Figure 1A**) was confirmed both by observing rapid AgRP neuron inhibition following chow presentation to overnight-fasted mice (**Figure 1B**) and by postmortem detection of GCaMP6s expression in AgRP neurons (**Figure 1A**). Either water or one of 4 doses of glucose (0.5, 1.0, 2.0, and 2.5 g/kg) was administered via the OG route in separate sessions. Monitoring commenced 25 min before OG administration and was maintained for 90 min thereafter, with dose order randomized across days. Prior to data analysis, CGM traces were processed with robust despiking, median filtering, and causal smoothing, then baseline-subtracted to yield the change of glucose over time (ΔG(t); see Methods, Glucose analysis).

**Figure 1.**
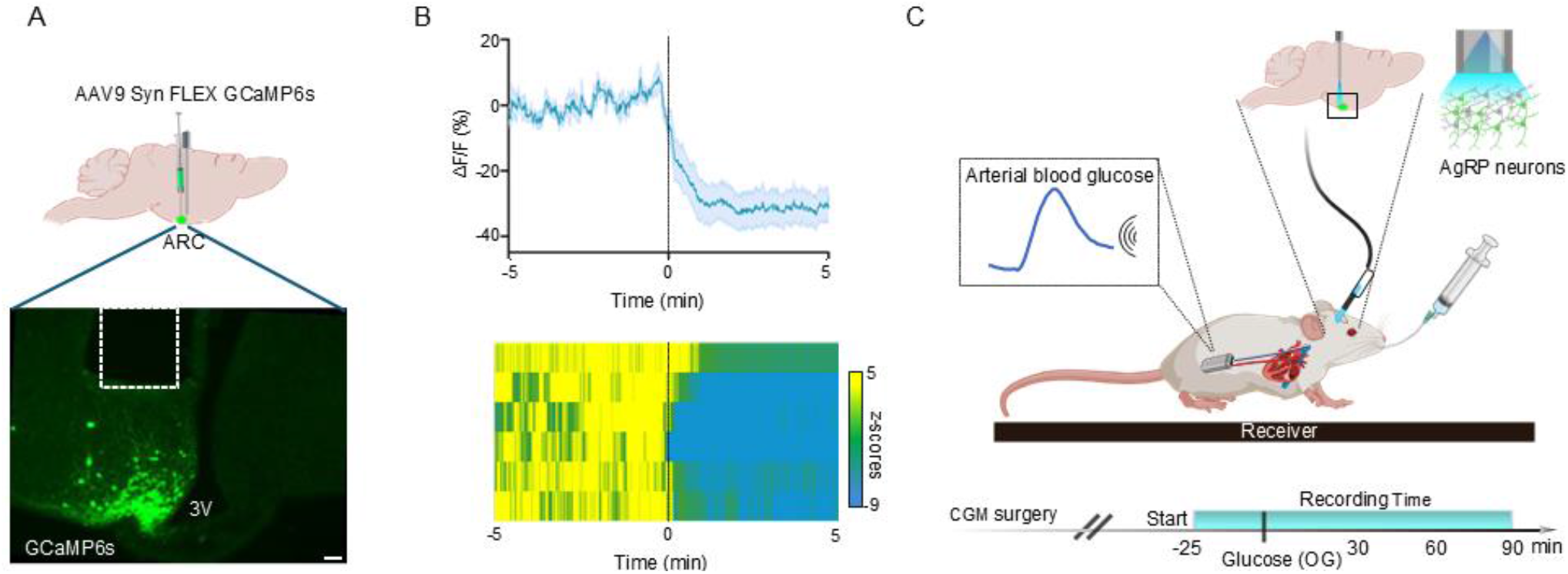
Experimental setup and validation of calcium activity response in AgRP neurons in vivo. (**A**) Schematic of viral injection and optic implant placement for photometry recordings of AgRP neuron activity (top), and representative image of GCaMP6s expression in the ARC (bottom; scale bar = 200 µm). (**B**) GCaMP6s responses to chow presentation (time = 0 min) in overnight-fasted AgRP-IRES-Cre mice (n = 6); mean ± SEM (top) and heat map of z-scored activity for individual mice (bottom). (**C**) Experimental design for simultaneous photometry and continuous glucose monitoring (CGM) in 5-hour fasted mice receiving glucose via oral gavage (OG).

As expected, oral glucose elicited dose-dependent increase in the BG level, with consistent temporal dynamics across sessions. Both the peak BG change from baseline (peak ΔG) and rate of BG rise increased in a dose-dependent manner (**Figure 2A-E**). Because the rate of BG decline after the peak was also proportional to the OG glucose dose (**Figure 2F**), overall recovery times were similar irrespective of dose. Consequently, while peak ΔG and the rates of rise and fall of BG were strongly dose-correlated (R^2^ = 0.38, 0.34, and 0.46, respectively), the incremental area under the blood glucose curve (iAUC) was not (R^2^ = 0.18) (**Figure 2D-G**).

**Figure 2.**
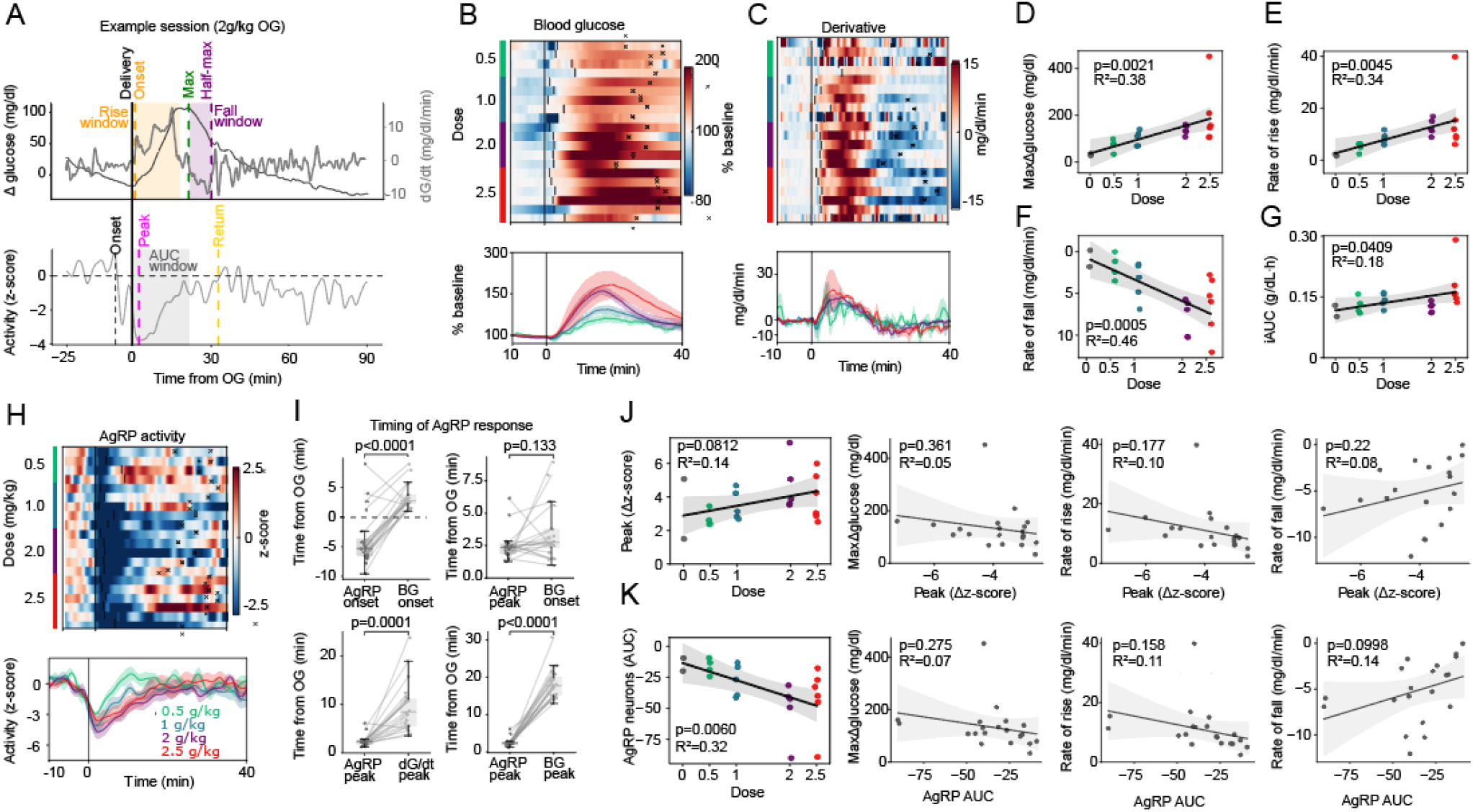
AgRP neuron responses to varying doses of oral glucose. (**A**) Example session showing blood glucose (BG) dynamics (**top**) and AgRP neuron activity (**bottom**) following OG of 2 g/kg glucose. (**B**) Heat-map of baseline-normalized BG levels for individual mice receiving different glucose doses (0.5–2.5 g/kg) over 40 minutes. Vertical lines mark the time of rise; x-marks indicate the time of return to half-maximum (**top**). The lower panel shows dose-averaged baseline-normalized BG. (**C**) Heat-map of BG derivatives after varying glucose doses, with time of rise and return to half-maximum indicated as in (**B, top**). The lower panel shows mean BG derivatives for each dose group. (**D–G**) Correlation between glucose dose and: peak ΔBG (**D**), BG rate of rise (**E**), BG rate of fall (**F**), and incremental area under the curve (iAUC) over 1 h after glucose or water administration (**G**). (n = 2– 6 per group). (**H**) Heat-map of AgRP neuron activity following different glucose doses, with rise onset and time of return to half-maximum indicated (top). The lower panel shows mean activity for each dose. (n = 4–6 per group). (**I**), Timing of AgRP neuron responses relative to BG dynamics: time of inhibition onset and inhibition peak compared to BG onset (top panels), and time of AgRP peak inhibition compared to time of peak BG derivative (bottom left) and peak BG (bottom right). (**J**) Relationship between peak AgRP inhibition (Δz-score, 0–10 min after OG) and glucose dose (first panel), as well as peak BG, BG rate of rise, and BG rate of fall (second–fourth panels). **K**, Relationship between negative AUC of AgRP responses (t = 0 min to peak BG) and glucose dose (first panel) or BG metrics (peak ΔBG, rate of rise, and rate of fall; second–fourth panels). Statistical comparisons between groups were performed using Wilcoxon tests (**J**), and linear relationships in (**D-G** and **J–K**) were assessed using ordinary least squares (OLS) regression with subject fixed-effects; shaded regions represent slope 95% confidence intervals.

### 2.2

Processing of z-scored FP signals from AgRP neurons in parallel with glucose traces (see Methods, Fiber photometry analysis) reveals that as expected [12,16], the activity of these neurons was markedly and rapidly decreased by OG glucose administration and remained suppressed for tens of minutes as the plasma BG levels rose and then subsequently declined (**Figure 2H**). Closer inspection of individual traces showed that in most cases, AgRP neuronal activity began to decline from its baseline *prior to* OG delivery itself, well before BG rise (AgRP onset –4.0±0.99 min (mean±SEM); N=22 sessions, *p* = 0.0006, Student’s 1-sample t-test). Indeed, AgRP neuron activity tended to reach peak inhibition around the time BG began to rise (AgRP peak +2.4±0.2 min vs BG rise +3.6±0.4 min (mean±SEM); *p* = 0.133, Wilcoxon signed-rank; **Figure 2I** top-right panel) and well before the BG rate-of-change had itself peaked (+2.4±0.2 vs peak dG/dt = 9.2±1.1 min, *p* = 0.0001, Wilcoxon signed-rank; **Figure 2I** bottom-left panel).

We then considered whether this earliest AgRP neuronal response (‘first phase’) reflects an anticipatory response conditioned by prior glucose exposures. In support of this hypothesis, the onset of AgRP neuronal inhibition occurred earlier among mice receiving their second glucose bolus compared to their first (**Supplementary Figure S1A**), suggesting a learned component. Additionally, an inhibition was detected when water was administered (instead of glucose) following prior glucose exposures (**Supplementary Figure S1B**). In contrast, this response was not observed when either water or glucose was given as the first OG administration (**Supplementary Figure S1C**). Thus, this ‘first phase’ decrease of AgRP neuronal activity appears to be a learned and anticipatory response, consistent with known inhibition of AgRP neurons to pre-ingestive cues such as food scent [13].

Peak AgRP neuronal inhibition following OG glucose coincided with the onset of rising BG levels (**Figure 2I**, right panel), but unexpectedly, its magnitude (captured by the peak negative z-score in a 0-10 min window following glucose administration) was invariant. Stated differently, peak inhibition did not vary with glucose dose, peak BG, or either the rate of rise or rate of fall of BG (**Figure 2J**), indicating a largely stereotyped, administration-aligned population response whose amplitude is independent of either the amount of oral glucose given or the subsequent glycemic response. Consistent with this early-phase dissociation, the broader suppression of AgRP neuron activity measured by incremental area under the curve (iAUC) from delivery (t=0) to time of peak plasma BG likewise showed no meaningful relationship to post-absorptive BG metrics (**Figure 2K**; see Supplement for iAUC analyses), despite varying with glucose dose.

### 2.3

To investigate the relationship between the glycemic excursion and AgRP neuron activity subsequent to its rapid initial decline (which includes the last phase of AgRP neuronal inhibition and its subsequent recovery towards its baseline, referred to herein as the ‘second phase’ response), we developed a model that isolates second phase activity from the first phase. (**Figure 3A**, see Methods, Early-response model).

**Figure 3.**
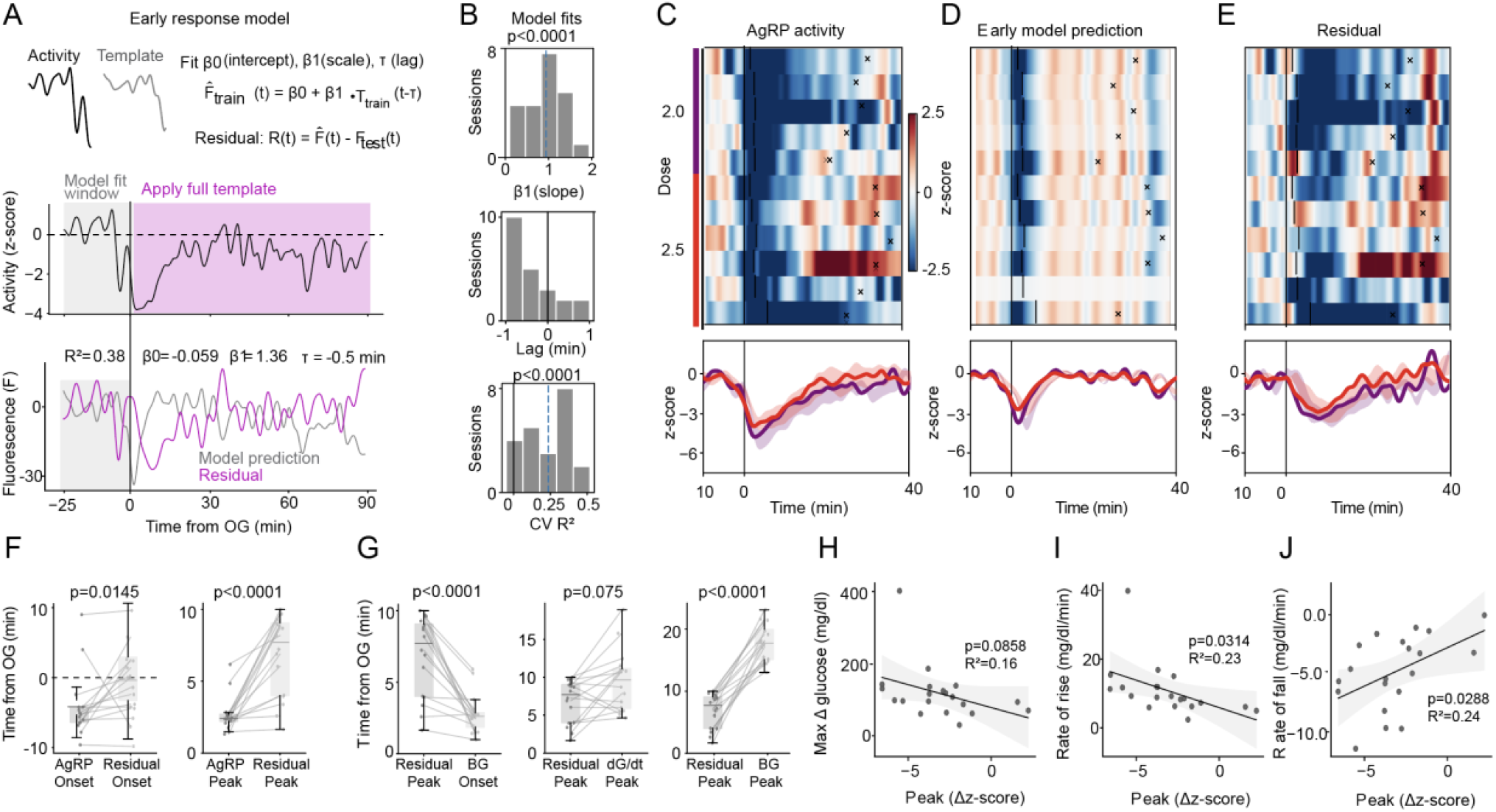
Dissocation between phase 1 and phase 2 responses using an early-response model. (**A**) Early-response model (see methods) illustrating the cross-validated prediction 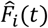 generated from low-dose (0 and 0.5 g/kg) sessions and the corresponding residual trace 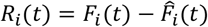, representing activity exceeding the early template (**top**). The middle panel shows AgRP neuron activity (z-score) used to fit the model window (−25 to +1 min) and the post-absorptive glucose template. The bottom panel displays the model prediction separated from the residual component. (**B**) Histograms showing early-response model fits across sessions: slope (top), lag (middle) and cross-validated R^2^ (bottom). (**C-E**) Heat-maps (top) and mean AgRP neuron activity (bottom) for the original (C), cross-validated prediction (**D**), and residual (**E**) traces in mice administered 2 or 2.5 g/kg glucose. (**F**) Timing of AgRP neuron responses: comparison of onset inhibition (**left**) and peak inhibition (**right**) before vs after model decomposition (residual). (**G**), Timing of AgRP neuron peak inhibition after correction: comparison with time of BG rise (left), BG peak derivative (middle), and BG peak (right). (**H-J**) Relationship between corrected AgRP neuron responses and BG metrics: peak ΔBG (**H**), BG rate of rise (**I**), and BG rate of fall (**J**). Statistical comparisons between groups were performed using the Wilcoxon signed-rank tests (**F** and **J**). Linear relationships in (**H–J**) were assessed with ordinary least squares (OLS) linear regression with fixed effects for subject; shaded regions indicate slope 95% CIs.

We fit the model’s lag and scale parameters using only the pre/early window (−20 to +1 min relative to glucose delivery) from glucose-administration sessions, optimizing these parameters within inner cross-validation folds (**Figure 3B**; see Methods, Early-response model). We then applied the resulting early-response prediction to a held-out low-dose test set (0 and 0.5 g/kg) and quantified deviations in the post-administration period (time points never used for fitting) as residuals, 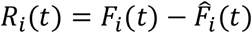. In this framework, residuals provide a session-by-session estimate of late activity not explained by the initial, BG-trajectory–independent, stimulus-locked component: the more strongly the second phase covaries with glycemic dynamics, the larger (more negative) the deviation from the early-model prediction.

Consistent with this decomposition, the early-response model closely captured the rapid initial decrease in AgRP neuronal activity, but it did not account for the sustained suppression that followed at higher glucose doses. Residuals became increasingly negative over time, peaking several minutes after BG began to rise, before gradually returning toward baseline (**Figure 3A**). Population heatmaps of the original, fitted, and residual traces confirmed this decomposition. Thus, the second phase component was weak or absent at low glucose doses but emerged prominently as extended inhibitory residual structure at higher doses (2 and 2.5 g/kg, **Figure 3C-E**).

We next interrogated the relationships between second phase AgRP neuronal activity and both the BG change over time and *the rate of BG change* (*i.e*., the BG derivative, which represents the direction of future BG levels). While the raw AgRP peak neuronal inhibitory response occurred concurrently with the initial rise in BG, this early response was captured by the early model, with peak inhibition in the residual now occurring later and aligning more closely with time of peak BG rise (**Figure 3F-G**). Accordingly, while peak residual inhibition occurred well before the BG peak, it no longer differed from the peak of the glucose derivative (**Figure 3G**). Interestingly, while the degree of maximum residual inhibition remained uncorrelated with the peak ΔG (**Figure 3H**), it correlated significantly with rates of BG rise and fall (R^2^ = 0.23 and R^2^ = 0.24, respectively; **Figure 3I-J**). These findings indicate that second-phase residual magnitude scales with the rate of glycemic change. By comparison, the same procedure in mice receiving either OG water or low-dose glucose (0.5 mg/kg) produced small residuals with no consistent stimulus-aligned structure and no significant relationship to glucose dynamics (**Supplementary Figure S2**).

### 2.4

To quantify how the two phases of the AgRP neuronal response are coupled to the change in glycemia, we used lagged cross-correlation [20]. Correlations were computed over ±20 min lags and evaluated against pseudosession nulls [21] (obtained by pairing FP with circularly shifted glucose traces; see Methods, Cross-correlation analysis). In the Early-response model FP signal, cross-correlations with BG and its derivative (based on an OG glucose dose of 2 mg/kg) were strong and skewed toward negative lags: the peak correlation occurred when AgRP neuron activity led the blood glucose signal by ~10–15 min and its derivative by ~0-10 min (**Figure 4A-B**), consistent with the first-phase response arising as an anticipatory signal.

**Figure 4.**
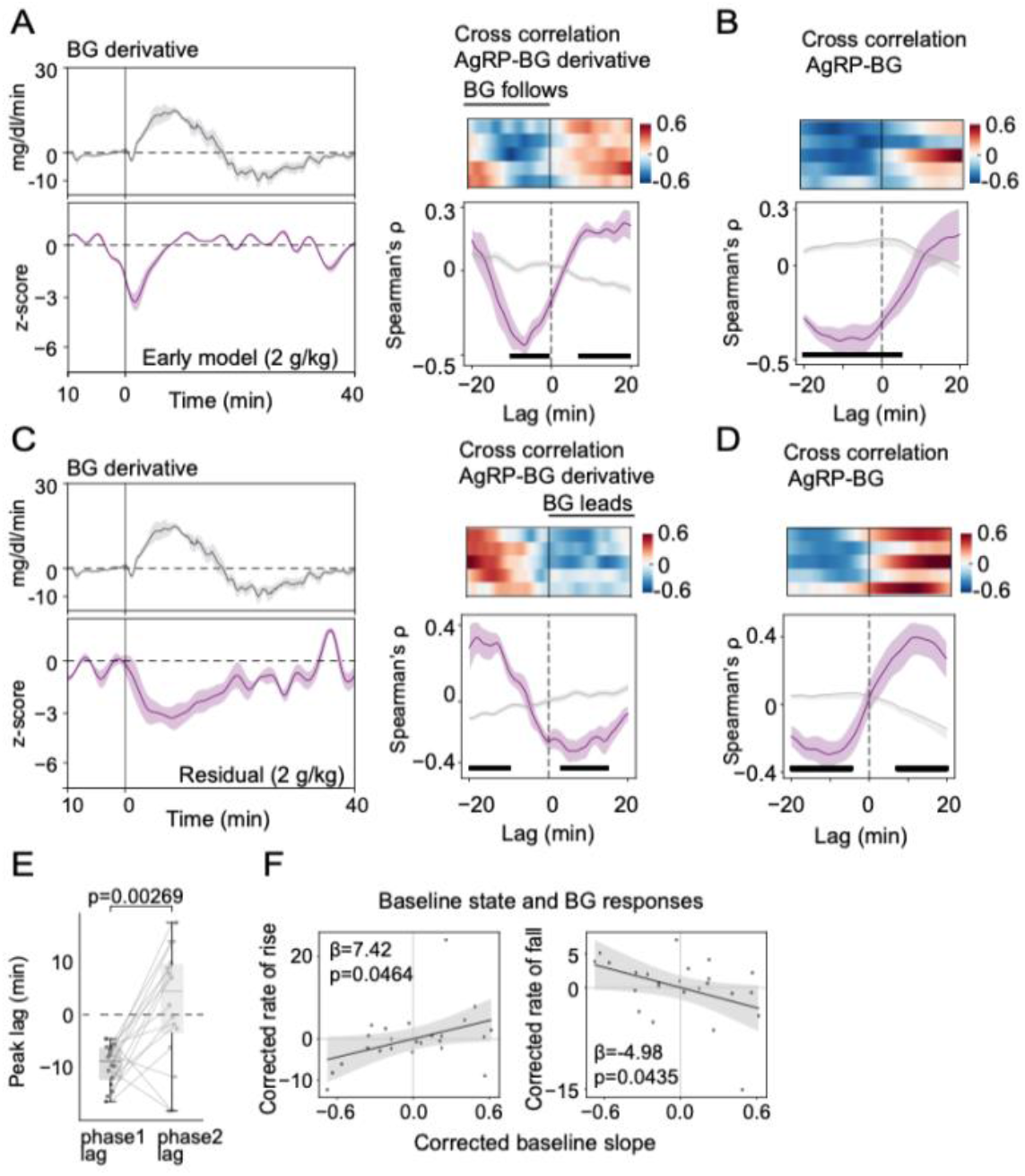
Temporal relationship between AgRP neuron activity and BG. (**A** and **B**) Mean BG derivative (**A, top**) and early-response modeled AgRP neuron activity (**A, bottom**, z-scores) after a 2 g/kg glucose dose. Lagged cross-correlation between the early AgRP response and the BG concentration (**A, right**) or BG derivative (**B**); vertical ticks indicate bins where *p*<0.01. (**C** and **D**) Mean BG derivative (**C, top**) and residual component of AgRP neuron activity after a 2 g/kg glucose dose (**C, bottom**). Lagged cross-correlation between the residual component and BG concentration (**C, right**) or BG derivative (**D**). Both FP and BG time series were rank-transformed (Spearman), and correlations were computed over ±20-min lags and evaluated against pseudosession-derived null distributions (Wilcoxon rank-sum tests with Benjamini-Hochberg correction for multiple comparisons; vertical ticks indicate bins where *p*<0.01). (**E**) Comparison between peak lag before and after correction (phase 1 vs phase 2). (**F**) Relationship between AgRP neuron baseline state and BG rate of rise (**left**) or BG rate of fall (**right**). Slope from linear mixed effects model with corrections for OG dose, subject, and baseline BG level.

After subtracting the early-response template from the raw FP signal, however, this coupling pattern changed. Residual activity remained significantly correlated with the blood glucose level, but with somewhat lower peak magnitude (**Figure 4C**). Importantly, the peak cross-correlation with the glucose derivative shifted to a short positive lag of ~5 min, with residual AgRP activity following rather than preceding the glucose derivative, and occurred later than the Early-model correlation (**Figure 4D-E**). This positive-lag trough significantly exceeded pseudosession nulls and was consistent across animals, indicating that once the early, administration-locked component is removed, the remaining activity primarily tracks the BG rate of change, rather than the BG level itself.

### 2.5

To quantify the extent to which baseline AgRP neuron state (prior to oral glucose delivery) predicts the subsequent glycemic response to a given glucose load, we analyzed a slope parameter generated from each session by the Early-response model. This parameter provides a measure of how strongly that session’s baseline and immediate post-gavage neuron activity conform to the average early template (β_1_ in **Figure 3A**). We interpreted this as a simple baseline “state” metric, where a larger slope indicates greater-than-average inhibition in the baseline and early-response periods, and vice versa.

Across sessions, greater slopes predicted increased rates of BG rate of rise and fall for a given glucose dose (**Figure 4F**). This finding implies that differences in pre-absorptive AgRP state covaries with trial-to-trial variability in the glycemic impact of the same nominal glucose load. A complementary baseline measure derived from the low-frequency structure of the FP signal during the pre-administration period showed a similar relationship to dose-corrected iAUC (**Supplementary Figure S3**), confirming that slow inhibition at baseline predicts a greater glycemic excursion with a more rapid rise and fall.

### 2.6

To assess whether AgRP neurons respond to changes in blood glucose independently of gut–brain signaling, we monitored their activity both before and after intraperitoneal (i.p.) injection of either saline (n = 3) or glucose (2g/kg; n = 5). As expected, AgRP neurons were inhibited following i.p. glucose administration relative to saline (glucose: z-score = −0.86% ± 0.38%; saline: z-score = +0.84% ± 0.44%, **Supplementary Figure S4**). Beyond this, however, two major differences were observed when compared to the response oral glucose. First, the anticipatory ‘first phase’ response seen with oral glucose was not observed. Second, the magnitude of AgRP neuron inhibition by IP glucose was small in comparison to oral glucose delivery, highlighting the important role played by gut-brain signals in the response. Together, these findings indicate that while AgRP neurons respond to changes in circulating glucose levels independently of gut–brain signaling, the latter plays a qualitatively more important role in the effect of glucose to inhibit these neurons.

## 3. Discussion

In addition to their potent stimulatory effects on food intake, AgRP neurons are increasingly recognized for their role in glucose homeostasis [22–27]. For example, these neurons are activated in most forms of diabetes [19,28–30], and in a mouse model of type 2 diabetes, AgRP neuron hyperactivity is required for hyperglycemia, but is dispensable for hyperphagia and obesity [17]. Conversely, the prolonged antidiabetic response to centrally administered fibroblast growth factor 1 (FGF1) is linked to sustained AgRP neuron inhibition [31]. Moreover, AgRP neurons have been described as “glucose-inhibited” based on *ex vivo* studies [32], and in vivo fiber photometry studies show that their activity decreases following either oral glucose or glucose infusion into the hepatic portal vein [16]. Together, these observations suggest a close relationship between AgRP neuron activity and glycemic state. Accordingly, we hypothesized that following an oral glucose load, AgRP neuron activity would exhibit temporally structured changes aligned with the glycemic response.

To test this hypothesis, we implemented a strategy wherein AgRP neuronal activity and the arterial glucose level are monitored continuously both prior to and after OG glucose administration. We find that in sessions conducted in mice previously subjected to OG glucose administration, reduced AgRP neuronal activity is detectable *before* that session’s OG administration. This early inhibitory response was consistently observed across subsequent sessions, regardless of whether water or glucose was administered. Thus, this ‘first phase’ response appears to be mounted in anticipation of an oral glucose load, a possibility strengthened by the finding that neither its timing nor its magnitude is impacted by glucose dose (which, of course, cannot be anticipated).

This anticipatory response is consistent with evidence that in fasted mice, rapid AgRP neuron inhibition is elicited merely by the sight or smell of food, even if the food is not eaten [11,13,14]. If food is not consumed, however, this rapid inhibitory response is both modest and short-lived when compared to the response to a consumed meal [13]. Furthermore, the neurocircuitry implicated in this anticipatory AgRP neuron response appears to be distinct from that underlying the more sustained inhibition induced by food ingestion. Specifically, AgRP neuron inhibition by sensory cues involves projections from the lateral hypothalamic area to inhibitory GABAergic neurons in the dorsomedial hypothalamus (DMH) that lie upstream of and inhibit AgRP neurons [33], whereas signals emanating from the GI tract are implicated in the sustained inhibitory response. Additional studies are warranted to determine whether the former pathway also underlies the anticipatory ‘first phase’ response reported here, and the extent to which this early response carries information about both the animal’s expectation and its internal state before ingested nutrients are absorbed.

That the early decrease of AgRP neuron activity *1)* precedes OG glucose delivery, *2)* appears to be anticipatory in nature, and *3)* is predictive of the glycemic excursion (*i.e*., greater early inhibition predicts better glucose tolerance) raises a series of questions. Is it possible, for example, that early AgRP neuron suppression participates in cephalic phase/autonomic control of glucose handling? Such responses can influence gastric secretions, gastric emptying, intestinal transit, insulin secretion, and hepatic glucose output, although the role played by AgRP neurons in these responses awaits further study. Consistent with this possibility, however, is evidence that chemogenetic AgRP neuron activation both impairs hepatic insulin sensitivity and increases hepatic glucose production [34], metabolic adaptations characteristic of the fasted state.

An alternative (and non-mutually exclusive) explanation is that early AgRP neuron suppression primarily reports internal state (e.g., hunger/arousal): sessions starting from a higher hunger-related AgRP neuron activity baseline may exhibit larger early inhibition and, independently, altered glucose absorption and clearance dynamics and associated metabolic responses. Thus, while early AgRP dynamics are predictive of glycemic responses, establishing whether this reflects a causal role for AgRP neurons versus shared upstream, state-dependent control will require targeted manipulations and direct measurements of intermediate physiological steps.

Following the ‘first phase’ inhibitory response and the subsequent transition period during which inhibition peaks, AgRP neuron activity gradually rises towards its baseline level. Unlike the first phase response, this ‘second phase’ is consistently associated with BG metrics. To optimize the analysis of this relationship, we constructed a model wherein the ‘first phase’ response is effectively subtracted to enable analysis of the second phase. This analysis showed that during the second phase, slower recovery of AgRP neuron activity (greater inhibition) is associated with more rapid rates of BG rise and fall. Interestingly, the second phase response dynamics aligned more closely with the BG *rate of change* (derivative) than with the absolute BG level. This observation parallels the finding that following an oral glucose load, hypothalamic orexin neuron activity and BG rate of change are tightly linked to one another [35]. Since the rate of change of a parameter [31] is more predictive of future state than its absolute value, sensory detection of the rate of change aligns with an ‘allostatic’ control mechanism that combines interoceptive processing with learned experience to mount anticipatory responses based on predictions about future state [36,37].

Importantly, however, our studies were not designed to test whether AgRP neurons directly sense and respond to changing blood glucose levels. Rather, the temporal coupling we observe likely involves indirect mechanisms, potentially including gut–brain signaling, portal vein glucose sensing, and/or vagal afferent pathways. Because our oral gavage paradigm preserves these physiological pathways while also allowing for anticipatory control, it enables characterization of the temporal relationship between AgRP neurons and blood glucose dynamics within an intact, biologically relevant context.

While it is conceivable that parenteral glucose (intraperitoneal or intravenous) administration might separate first and second phase responses from each other, we believe that this approach is not well suited to this purpose. At issue is that parenteral administration bypasses key gastrointestinal and portal signaling pathways while also producing glucose kinetics that differ substantially from physiological oral glucose absorption. Consistent with this interpretation, we report that glucose delivered IP elicits only a brief and modest inhibition of AgRP neurons [12], rather than the sustained second-phase component observed after oral glucose administration. Conversely, glucose infusion directly into a duodenal catheter (data not shown) elicits an AgRP neuron response resembling the phase 2 response reported here, but without an early anticipatory phase (phase 1), as expected since the animals are unaware that glucose is being administered [16]. Published evidence confirms this interpretation, as glucose delivered via gastric catheter or directly into the hepatic portal vein also suppresses AgRP neurons in a manner resembling the second-phase response, but without a preceding anticipatory component [12,16].

Together, these observations support the interpretation that second-phase dynamics primarily reflect gut- and portal-mediated signals rather than any direct sensing of circulating glucose. Nevertheless, it is interesting that the magnitude of the second phase response is coupled to the dynamics of the glycemic response. Future studies combining fiber photometry, CGM, and direct intragastric or intraduodenal glucose delivery will be important to more precisely dissect the relative contributions of anticipatory and post-ingestive signals to the AgRP neuron responses, and whether and how these changes of AgRP neuron activity impact the glycemic response.

In conclusion, we report that the AgRP neuron response to oral glucose comprises two functionally distinct phases, each with their own unique relationship to the glycemic response. The first phase, stimulus-aligned component (captured by a simple, BG-trajectory–independent template) predicts how a given glucose load will impact the subsequent glycemic response. While additional studies are needed, we interpret these findings to suggest that AgRP neurons help to transduce anticipatory information about impending nutrient delivery into adaptive metabolic responses that optimize glucose handling. Subsequent to this first phase is a second phase that *1)* remains after template subtraction, *2)* scales with dose-dependent BG metrics, and *3)* is coupled to BG dynamics – in particular, the rate of BG change. These findings lay the foundation for future investigation into whether a similar relationship exists between AgRP neurons and other circulating macronutrients, and how these relationships might be altered in metabolic disorders such as obesity and diabetes.

## Methods

### 4.1 Animal Studies

All animal experiments were conducted in accordance with the National Institutes of Health Guide for the Care and Use of Laboratory Animals, and all procedures were approved by the Institutional Animal Care and Use Committee at the University of Washington, Seattle, WA. AgRP-IRES-Cre mice (kindly provided to us by Dr. Streamson Chua, Jr Albert Einstein College of Medicine)) have previously been described [38] and are readily available from Jackson Laboratories (012899, Bar Harbor, ME, USA). Mice were individually housed in a temperature-controlled room on a 14:10 light:dark cycle under specific pathogen-free conditions with ad libitum access to drinking water and standard laboratory chow (PMI Nutrition, St. Louis, MO). Before experimental sessions, mice were fasted for either 16 h (for validation studies) or for 5 h (for analysis of the relationship between AgRP neuron activity and the BG level), as detailed in the text and figure legends.

### 4.2 Surgeries

To measure AgRP neuron activity, we stereotaxically injected unilaterally into the ARC (anterior-posterior (AP), −1.5 mm; mediolateral (ML), 0.3 mm; dorsoventral (DV, from dura), −5.85 mm, 400 nl, at a rate of 100 nl/min) an AAV containing a cre-inducible GCaMP6s cassette: pAAV9-CAG-FLEX-GCaMP6s (Addgene, Plasmid no. 100842, Watertown, MA, USA, titer 1×10^13^ per ml) into AgRP-IRES-Cre mice as previously described [39,40]. In the same surgery, a photometry cannula (MFC_400/430–0.48_6.1 mm_MF1.25_FLT, Doric Lenses, Franquet, Quebec, Canada) was implanted above the injection site (AP = −1.5 mm, ML = −0.3 mm, DV = −5.8 mm), as described [40,41]. Mice were allowed 2–3 wk for viral expression and recovery before FP recording sessions. Accurate viral targeting and fiber placement were verified by measuring AgRP neuron responses to chow refeeding after overnight fasting. Animals showing less than ΔF/F 20% response to chow were excluded from further experiments. Accurate targeting was also confirmed at the end of each experiment by histochemical detection of GCaMP6s and optic fiber placement.

Glucose sensors (Data Sciences International (DSI), St. Paul, MN, USA) were implanted through the left carotid artery of AgRP-IRES-Cre mice following the manufacturer’s guidelines and as previously described [20,42] with support of the NIDDK-funded Diabetes Research Center Metabolic and Cellular Phenotyping Core at the University of Washington. Mice were allowed 1 wk for recovery from surgery.

### 4.3 Fiber photometry

AgRP neuron activity was measured using a photometry processor (RZ 10X, Tucker-Davis Technologies (TDT), Alachua, Florida, USA) as previously described [20,39,40]. Briefly, two excitation LEDs were used for this study, 405 nm at 210 Hz and 465 nm at 330 Hz, power of 30-50 μW at the implant fiber tip. These LEDs were delivered to a filtered minicube (Doric Lenses, Franquet, Quebec, Canada) and passed through patch cords (MFP_400/430/1100– 0.57_0.45m_FCM-MF1.25_LAF, Doric Lenses, Franquet, Quebec, Canada) to the mouse photometry cannulae. The 405-nm LED was used as an isosbestic signal to control for movement and photobleaching. Digital signals were collected through the processor (RZ 10X, TDT), and data were captured using the software Synapse (TDT) and analyzed using custom Python code.

### 4.4 Continuous arterial glucose monitoring

HD-XG telemetry system (DSI, St. Paul, MN, USA) was used to continuously monitor blood glucose concentrations. Calibration of the implants was performed by OGTT test following the manufacturer’s instructions. CGM data was acquired by Ponemah software (DSI). For data synchronization, Synapse software generated TTL pulses that were sent to Ponemah via a signal interface box (DSI), allowing alignment of FP and CGM data acquisition. Data was sampled at 1 Hz.

### 4.5 Study protocols

Starting at the beginning of the light cycle, mice were subjected to a 5-h fast while remaining in their home cages. They were then connected to a patch cord for FP recordings and their home cage was placed on a receiver plate (DSI) for CGM. After a 20-min acclimation period, simultaneous FP and CGM recordings were initiated. The protocol began with baseline recordings for 25 min prior to OG administration of either water or one of four doses of glucose (0.5, 1, 2, and 2.5 g/kg). Each animal received each intervention in randomized order, and recordings continued for an additional 90 min post-gavage. For data analysis, the time of OG administration was defined as zero.

For intraperitoneal (i.p.) injection experiments, a separate cohort of 5-h-fasted mice was connected to a patch cord for FP recordings. After a 10-min baseline period, mice received an i.p. injection of either glucose (2 g/kg) or saline, and FP recordings continued following injection.

### 4.6 Data and statistical analysis

#### Glucose analysis: Pre-processing (filtering, smoothing, baseline-subtracting)

We first cleaned each glucose trace to remove artifacts while preserving timing. Noise was flagged using a robust median absolute deviation (MAD) rule (|y−median| > *k*·MAD with *k* = 2.0) and replaced by linear interpolation. We then applied a short centered median filter (7 samples = 7 s) to remove any remaining impulsive noise, followed by a causal moving average (60 samples = 60 s) that smooths high-frequency jitter yet keeps the response aligned in time. Finally, we defined a stable baseline from the pre-event period (−25 to 0 min from glucose administration) and worked with the baseline-subtracted signal, *ΔG*(*t*), so amplitudes are comparable across sessions.

To quantify dynamics, we computed a finite-difference derivative at 1-s spacing and smoothed it with the same 60-sample causal window. We defined response onset as the first sustained moment after t = 0 (time of OG) when either the derivative rose above a robust baseline threshold or the level cleared a baseline-scaled threshold, with a ≥1 min sustain to avoid false starts (see below). We located the peak level of *ΔG*(*t*) within 0–90 min and recorded the peak derivative in the early window (0–30 min after glucose delivery) to capture the time and level of sharpest rise. Recovery was summarized by the half-max time (first crossing back to the baseline + 50% of peak, computed on the raw series with linear interpolation). We defined a plateau phase where the smoothed derivative dropped into a near-zero band scaled to the preceding rise, then identified the return to baseline either by amplitude (first *ΔG*(*t*) ≤ 0) or by a sustained relaxation of the derivative back above a positive threshold. Overall exposure was captured by a positive-only incremental area under the curve (iAUC) over the entire post-delivery period (0–90 min), ensuring that only excursions above baseline contributed.

The first sustained rise in the BG level after *t* = 0 was identified by testing two independent conditions against robust, baseline-derived thresholds and requiring ≥1 min of continuous evidence (1 Hz → 60 consecutive samples). First, we smoothed the baseline-subtracted glucose *y*_0_(*t*) = *G*(*t*) − *median*_[−5,0]_*G* with a 2-min causal boxcar (≈120 samples at 1 Hz) and took a finite-difference derivative *dy*/*dt*. On the baseline segment (t < 0), we computed robust centers and scales:

- **Derivative (slope) threshold**. Let *d*_*pre*_be baseline derivatives. With median *m*_*d*_ and scaled MAD 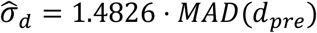, the slope threshold is

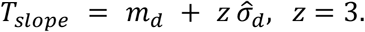
- **Level (amplitude) threshold**. Let *y*_0,*pre*_ be baseline values of *y*_0_. With scaled MAD 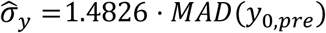, the level threshold is

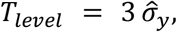

We then mark samples *t* ≥ 0 where either 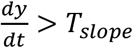 or *y* (*t*) > *T*_*level*_. Using a sliding window, we require this condition to hold for a continuous 1-min run before declaring onset; the onset time *θ* is the first sample meeting that sustain criterion.

### 4.7 Fiber photometry analysis

#### Post-processing

Photometry data were analyzed using custom Python scripts (https://github.com/nikhayes/fibphoflow). We removed shared motion/bleaching components from raw fluorescence data by regressing a reference (isosbestic) or drift proxy onto the signal channel and subtracting the fitted component. This targets very slow, non-neural trends so the remaining signal reflects neural activity more faithfully.

### Pre-processing (median, causal smoothing, low-pass, z-score)

Photometry signals were preprocessed with MAD-based de-spiking (k = 2.0), a centered median filter (7 samples) to eliminate outliers, and a causal moving average (60 samples) so responses remained stimulus-aligned. To amplify slow, stimulus-evoked responses we applied a zero-phase Butterworth low-pass filter defined by period (cutoff 4 min, order 3). For comparability across animals, we z-scored each trace within the baseline (−25 to 0 min), yielding *Z*_*FP*_ (*t*) with mean 0 and unit SD at baseline.

### Response metrics

#### Onset of low-frequency activity

We quantified response timing by tracking when low-frequency photometry power arose around glucose delivery. For each session, we computed a spectrogram of the baseline-z-scored trace *Z*_*FP*_ (*t*) using 20-min windows with 99% overlap. This yielded power spectral density *S*(*f, τ*) in cycles per minute (cpm) as a function of time *τ* relative to the event. We focused on a single low-frequency band corresponding to periods between 14 and 20 minutes,

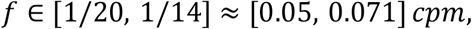

and retained only frequencies in this range. Band-limited power at each time bin was defined as

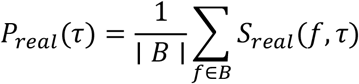

where *B* indexes the 14–20 min band. To test whether this power exceeded chance within the same session, we constructed a jittered null using pseudo-events. We drew *N* ≈ 200 pseudo-onset times, excluding a 5-min guard around the true glucose time. For each pseudo-event, we repeated the same segmentation and band-limited spectrogram, giving null band-power traces 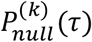. At each time bin we then computed an upper-tailed, rank-based p-value:

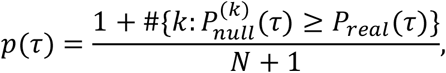

so that small *p*(*τ*) indicates that observed band power exceeds the null. We restricted detection to the symmetric window from −10 to +10 min around glucose delivery. The frequency-specific response onset time was defined as the first-time bin *τ*^∗^ ∈ [−10,10] where band power significantly exceeded the null. Sessions without any bin meeting this criterion were classified as having no detectable response in the 14–20 min (≈0.05–0.071 cpm) band.

#### Baseline low-frequency power (PSD slope)

Within the baseline window we estimate Welch’s power spectral density (PSD) and fit a simple aperiodic 1/*f*^*k*^ model in log–log space. The slope *k* summarizes bias towards slow baseline structure (higher slope □ greater low-frequency bias). *Peak z-response*. In an early window (0–15 min) we report the maximal *Z*_*FP*_. This captures the sharpest early deflection relative to baseline variability.

### 4.8 Area under the curve (AUC)

We compute an incremental AUC from glucose delivery (t = 0) to the time of peak blood glucose using NumPy’s trapezoidal rule, after setting all values above baseline to 0; this integral therefore reflects only negative deflections. We also compute a total AUC from t = 0 to the end of the session using the unmodified signal (no clipping), providing an overall measure of deviation from baseline.

### 4.9 Early-response model

To isolate stereotyped early, dose-agnostic dynamics from putative feedback components, we fit a simple lagged linear model only on pre-response activity and then examine residuals predicted from held-out sessions.

#### Model and variables

Let *F*_*i*_(*t*) be the z-scored FP for session *i*, and let *F*_*train*_ (*t*) be a template formed by averaging the baseline-aligned FP across the *training* sessions (see below). For a candidate lag *τ* ∈ [−1: 0.25: 1] minutes, we fit

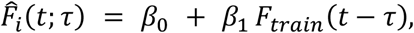

with *β*_0_ (intercept), *β*_1_ (slope/scale), and *τ* (temporal offset). Parameters *β*_0_, *β*_1_are estimated by ordinary least squares for each *τ*; *τ* is chosen by inner-fold cross-validation (below). This “template-regression” captures how strongly each session follows the average early pattern, while allowing a small temporal shift.

#### Training set, template construction, and test set

Training data comprised oral glucose sessions at 1.0, 2.0, and 2.5 mg/kg. For each session, we extracted the pre-peak window (−25-+1 min relative to the event) and resampled on the native 1 Hz grid. The template *F*ˉ_*train*_ (*t*) was computed as the across-session median at each time point to be robust to outliers. The test set consisted of water and 0.5 mg/kg sessions, held out entirely during model selection.

#### Inner-fold cross-validation (model selection)

We performed K-fold cross-validation (CV) across *training sessions* (folds stratified by subject when possible). For each candidate lag *τ* (e.g., a small grid around zero), we:

1. Rebuilt 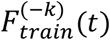 from the K−1 training folds (leaving fold *k* out).
2. Fit *β*_0_, *β*_1_on those folds by minimizing squared error on *t* ∈ [−25, +1]min.
3. Predicted the left-out fold using 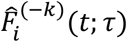 and accumulated validation error.

We selected *τ*^∗^ that minimized average validation mean-squared error (MSE) (tie-break by higher *R*^2^), then refit *β*_0_, *β*_1_on all training sessions using the template built from all training data and the chosen lag *τ*^∗^.

#### Evaluation on held-out sessions and residuals

With *τ*^∗^ and (*β*_0_, *β*_1_) fixed, we generated predictions 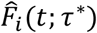 for each session from held-out averaged activity on water/0.5 mg/kg sessions. We then computed residual FP

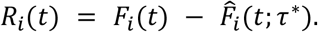

Because the model is fit exclusively on pre-peak-response activity and predictions are generated from low-dose glucose sessions, these residuals emphasize deviations that cannot be explained by the dose-agnostic early pattern that dominates low-dose glucose responses, including feedback-like components or dose-sensitive dynamics. For summary displays and statistics, we aggregated residuals over analysis windows (e.g., 0–10 min early, 10–60 min late) and, when needed, averaged across held-out sessions to visualize the typical “beyond-early-phase” deviation profile.

### 4.10 Cross-correlation analysis (Spearman, nulls, aggregation)

To characterize FP–glucose coupling and residual FP-glucose coupling without assuming linearity, we rank-transformed both series (Spearman) and computed lagged correlations over ±20 min in 1 s steps. We assessed specificity with pseudosession nulls generated by pairing FP with circular shifts of other sessions’ glucose (with a minimum 20 min offset). For group summaries, we Fisher-transformed per-session correlations, averaged in z-space (which stabilizes variance), mapped back to ρ for plotting, and controlled multiple comparisons across lags using a Benjamini-Hochberg correction for multiple comparisons.

### 4.11 Statistics

All statistical tests were two-sided. Statistical analyses were performed in Python (scipy.stats) and Prism 8.0 (GraphPad Software). The experimental unit for all analyses was the subject. Paired two-group comparisons were performed using the Wilcoxon signed-rank test, and unpaired two-group comparisons were performed using the Wilcoxon rank-sum test (Mann–Whitney U test). Linear relationships between continuous variables were evaluated using ordinary least squares (OLS) regression, with fixed effects for subject and, where indicated, baseline blood glucose and dose. OLS models were fit with robust standard errors. Statistical significance was defined as α = 0.05. For lagged cross-correlation analyses, p values were corrected for multiple comparisons across all lags within an analysis using the Benjamini–Hochberg false discovery rate procedure. Data are reported as median and interquartile range for box-and-whisker plots, and as mean ± SEM for line graphs. Outlier removal for continuous data is described in the data preprocessing section of the Methods. Exact p values are reported in the figures and figure legends.

### 4.12 Implementation notes

All data were sampled at 1 Hz. All analyses were implemented in Python using NumPy/Pandas for array and table operations, SciPy for filtering and frequency-based analysis (Savitzky–Golay, Butterworth/filtfilt, Welch PSD, spectrogram), scikit-learn for linear modeling, and Matplotlib for figures. All configurable parameters (windows, orders, thresholds, lags, null counts) are exposed via MetricsConfig (glucose) and FPConfig (photometry) and are reported in methods and accompanying code to support full replication.

## AUTHORSHIP NOTE

MG and AB are co-first authors

## ACKNOWLEDGMENT

The authors appreciate the technical assistance provided by Dr. Michael Bruchas, University of Washington, and Lisa Beutler and Nikolas Hayes (Northwestern University) for in vivo fiber photometry studies and analysis including custom scripts, and mouse colony management by Vincent Damian at the University of Washington.

## AUTHORSHIP CONTRIBUTION STATEMENT

**Micaela Glat:** Writing review & editing, Writing original draft, Supervision, Resources, Project administration, Methodology, Investigation, Formal analysis, Data curation, Conceptualization. **Anna Bowen:** Writing review & editing, Writing original draft, Supervision, Resources, Project administration, Methodology, Investigation, Funding acquisition, Formal analysis, Data curation, Conceptualization. **Yang Gou:** Methodology, Investigation, Data curation, Conceptualization. **Elizabeth Giering:** Methodology, Investigation. **Jarrad M. Scarlett:** Writing review & editing, Writing original draft, Resources, Project administration, Conceptualization. **Gregory J. Morton:** Writing review & editing, Writing original draft, Resources, Project administration, Conceptualization. **Michael W. Schwartz:** Writing review & editing, Writing original draft, Supervision, Resources, Project administration, Methodology, Investigation, Formal analysis, Data curation, Conceptualization.

## DECLARATION OF COMPETING INTERESTS

The authors declare the following financial interests/personal relationships which may be considered as potential competing interests: Michael W. Schwartz reports financial support was provided by National Institute of Diabetes and Digestive and Kidney Diseases. Jarrad Scarlett reports financial support provided by National Institute of Diabetes and Digestive and Kidney Diseases and the US Department of Defense. Gregory Morton reports financial support was provided by National Institute of Diabetes and Digestive and Kidney Diseases. Michael Schwartz reports a relationship with Novo Nordisk Inc that includes: funding grants.

## FUNDING

This work was supported by grants to: M.W.S. (NIDDK Grants DK083042 and DK101997), Y.G. (Cystic Fibrosis Foundation SINGH19R0 and SINGH24R0), J.M.S. (NIDDK Grants K08 DK114474, R03 DK128383, and DoD W81XWH2110635), G.J.M. (R01 DK089056, and R01 DK124238), A.B (Howard Hughes Medical Institute Hanna H. Gray Fellowship), the NIDDK-funded Nutrition Obesity Research Center (DK035816), and the NIDDK-funded Diabetes Research Center (DK017047) at the University of Washington. Funding in support of these studies was also provided to M.W.S. through an agreement with Novo Nordisk (CMS-431104).

## DATA AND CODE AVAILABILITY

All data are available at: https://doi.org/10.6084/m9.figshare.31891579. Companion code is available at: https://github.com/thebowenlab/glucose-photometry.git

## Supplemental figures

**Supplementary Figure S1.**
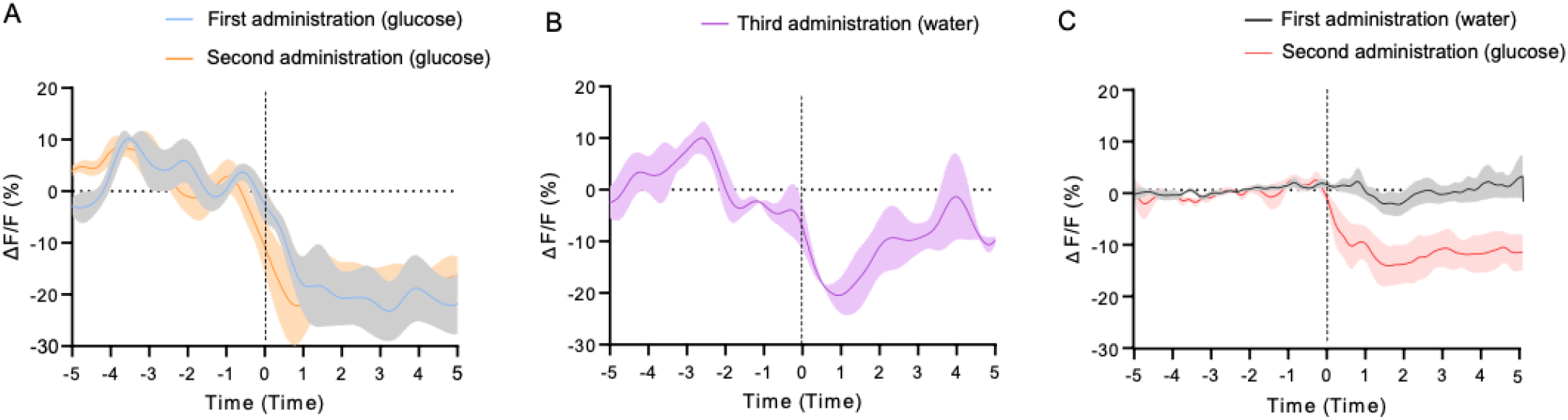
AgRP neurons exhibit anticipatory responses to glucose administration via OG. (A) Mean AgRP neuron responses to the first and second OG glucose (2g/kg) (n = 3). (B) Mean AgRP neuron activity when water was administered after prior glucose exposures (n = 3). (C) AgRP neuron activity when water was given as the first OG, followed by 2.0 g/kg glucose on the subsequent gavage (n = 1).

**Supplementary Figure S2.**
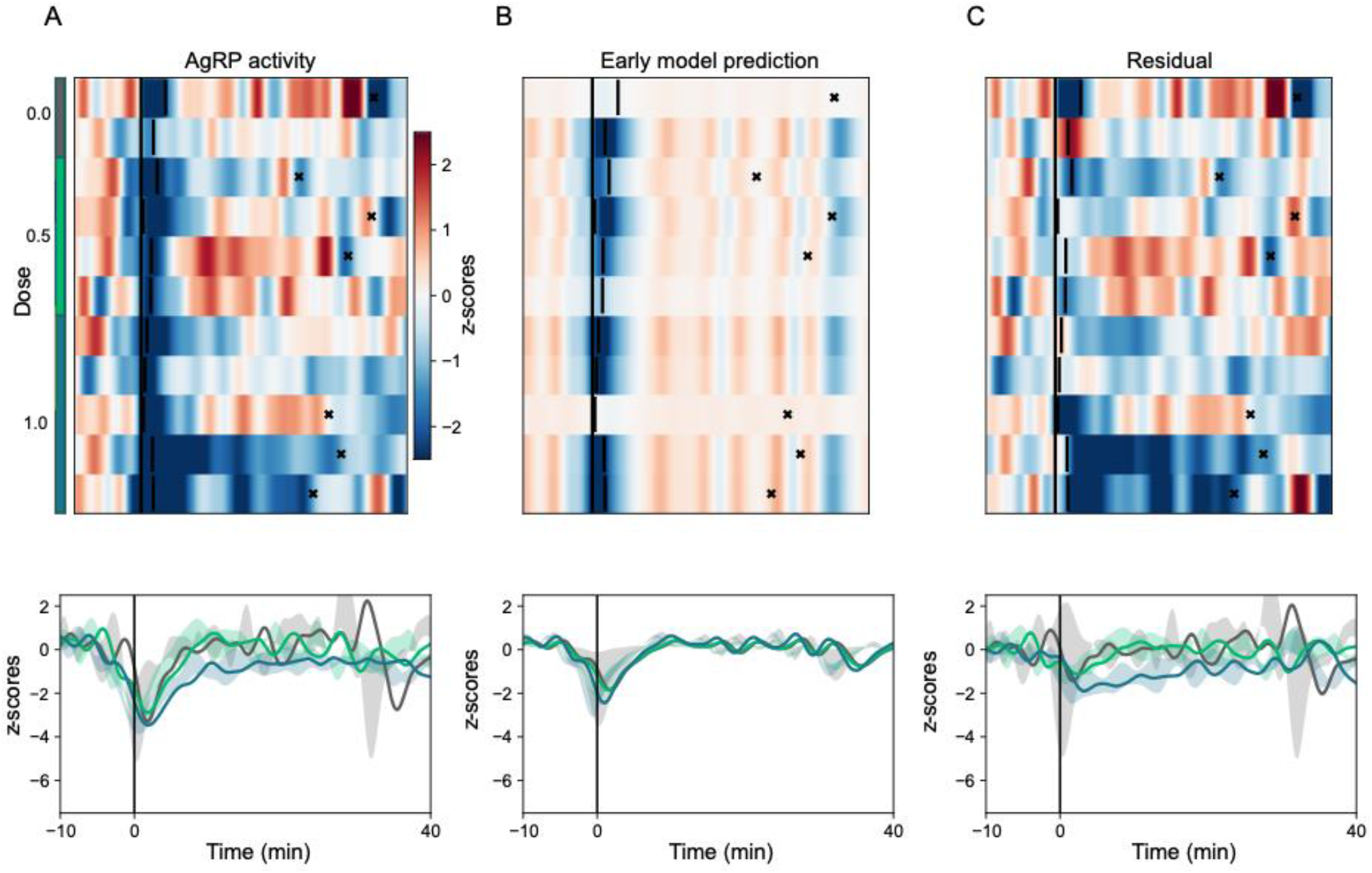
Dissociation between phase 1 and phase 2 responses using an early-response model in low doses of glucose. (A-C), Heat-maps (top) and mean AgRP neuron activity (bottom) for the original (A), fitted (B), and residual (C) traces in mice administered with water (0.0), 0.5, or 1.0 g/kg glucose.

**Supplementary Figure S3.**
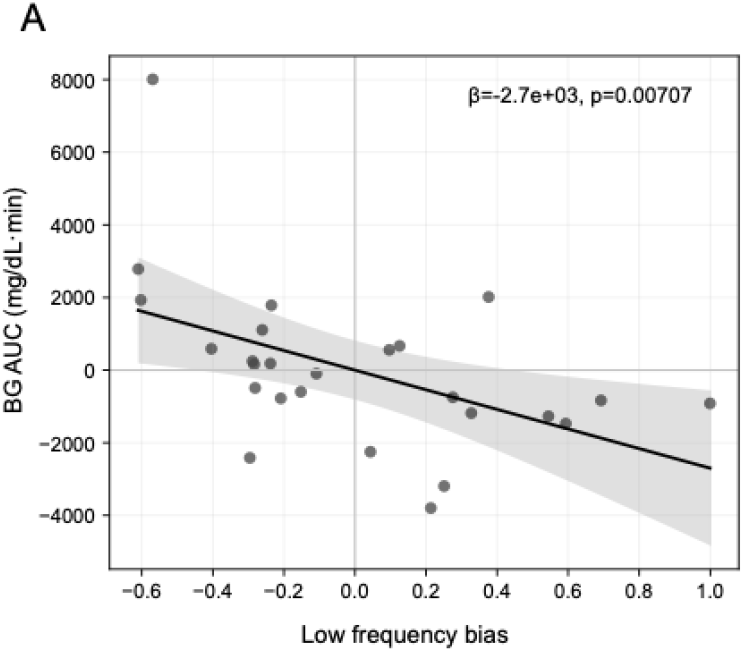
Relationship between baseline low-frequency power and subsequent BG dynamics. (A), Relationship between individual session aperiodic slope in AgRP neuron activity (higher slope → greater low-frequency bias in baseline period) and incremental AUC of BG level. Slope from linear mixed effects model with corrections for OG dose and subject.

**Supplementary Figure S4.**
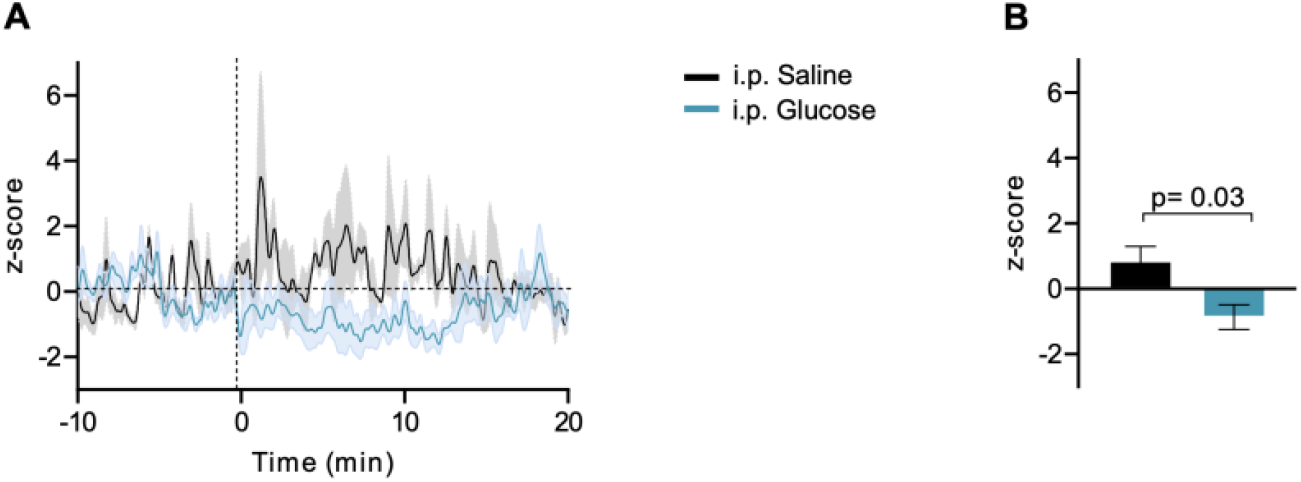
AgRP neuron inhibition following i.p. injection of glucose (2g/kg) or saline. (A) GCaMP6s responses to i.p. glucose (2g/kg) or saline in 5-hour fasted mice. (B) Quantification of z-score from (A) over the 0-15 min period following injection. Mann-Whitney U, p=0.03, glucose: n = 5; saline: n=3.

## Notes

Conflict of interest: Funding in support of these studies was provided to M.W.S. through an agreement with Novo Nordisk (CMS-431104).

https://doi.org/10.6084/m9.figshare.31891579

https://github.com/thebowenlab/glucose-photometry.git

### Competing Interest Statement

Funding in support of these studies was provided to M.W.S. through an agreement with Novo Nordisk (CMS-431104).

### Summary of Updates

In this revision the introduction and discussion were revised to add emphasis to discussion of prior work identifying predictive responses of AgRP neurons to food cues; the discussion was revised to clarify likely sources of second phase activity; typos were corrected in axes and panel labels in figures 2 and 3; a new study and supplemental figure S4 was added to examine AgRP neuron response to IP glucose administration (to isolate the putative humoral component of second phase response); the methods section was lightly revised to improve readability; a statistical analysis section was added to the methods.

